# Temporal recording of mammalian development and precancer

**DOI:** 10.1101/2023.12.18.572260

**Authors:** Mirazul Islam, Yilin Yang, Alan J. Simmons, Vishal M. Shah, Musale Krushna Pavan, Yanwen Xu, Naila Tasneem, Zhengyi Chen, Linh T. Trinh, Paola Molina, Marisol A. Ramirez-Solano, Iannish Sadien, Jinzhuang Dou, Ken Chen, Mark A. Magnuson, Jeffrey C. Rathmell, Ian G. Macara, Douglas Winton, Qi Liu, Hamim Zafar, Reza Kalhor, George M. Church, Martha J. Shrubsole, Robert J. Coffey, Ken S. Lau

**Affiliations:** Epithelial Biology Center, Vanderbilt University Medical Center, Nashville, TN, USA; Department of Cell and Developmental Biology, Vanderbilt University, Nashville, TN, USA; Department of Computer Science and Engineering, Indian Institute of Technology Kanpur, Kanpur 208016, India; Department of Biological Sciences and Bioengineering, Indian Institute of Technology Kanpur, Kanpur 208016, India; Program in Chemical and Physical Biology, Vanderbilt University School of Medicine, Nashville, TN, USA; Center for Stem Cell Biology; Department of Molecular Physiology and Biophysics; Vanderbilt University, Nashville, TN 37232, USA; Department of Biostatistics and Center for Quantitative Sciences, Vanderbilt University Medical Center, Nashville, TN, USA; Cancer Research UK Cambridge Institute, University of Cambridge, Li Ka Shing Centre, Robinson Way, Cambridge CB2 0RE, UK; Department of Bioinformatics and Computational Biology, The University of Texas MD Anderson Cancer Center, Houston, TX, USA; Vanderbilt Center for Immunobiology, Department of Pathology, Microbiology, and Immunology, Vanderbilt University Medical Center, Nashville, TN 37232, USA; Department of Biomedical Engineering, Department of Genetic Medicine, Johns Hopkins University School of Medicine, Baltimore, MD 21205, USA; Department of Genetics, Harvard Medical School, Boston, MA, USA; Wyss Institute for Biologically Inspired Engineering at Harvard University, Boston, MA, USA; Department of Medicine, Division of Epidemiology, Vanderbilt University Medical Center, Nashville, TN, USA; Department of Surgery, Vanderbilt University Medical Center, Nashville, TN, USA; Vanderbilt-Ingram Cancer Center, Vanderbilt University Medical Center, Nashville, TN, USA

## Abstract

Key to understanding many biological phenomena is knowing the temporal ordering of cellular events, which often require continuous direct observations [1, 2]. An alternative solution involves the utilization of irreversible genetic changes, such as naturally occurring mutations, to create indelible markers that enables retrospective temporal ordering [3-8]. Using NSC-seq, a newly designed and validated multi-purpose single-cell CRISPR platform, we developed a molecular clock approach to record the timing of cellular events and clonality *in vivo*, while incorporating assigned cell state and lineage information. Using this approach, we uncovered precise timing of tissue-specific cell expansion during murine embryonic development and identified new intestinal epithelial progenitor states by their unique genetic histories. NSC-seq analysis of murine adenomas and single-cell multi-omic profiling of human precancers as part of the Human Tumor Atlas Network (HTAN), including 116 scRNA-seq datasets and clonal analysis of 418 human polyps, demonstrated the occurrence of polyancestral initiation in 15-30% of colonic precancers, revealing their origins from multiple normal founders. Thus, our multimodal framework augments existing single-cell analyses and lays the foundation for *in vivo* multimodal recording, enabling the tracking of lineage and temporal events during development and tumorigenesis.

## Introduction

Mammalian development originating from a fertilized egg (zygote) is a remarkable process, comprising a highly orchestrated series of cell divisions and lineage diversifications [9]. The classic reconstruction of the *C. elegans* cell lineage and temporal histories from the zygote stage is a significant milestone for the field of developmental biology [10, 11]. Tumorigenesis shares cellular and molecular events with embryonic development, many of which have been recently appreciated [12-16]. The molecular mechanisms underpinning these events remain central questions in cancer biology. Fundamental to understanding these mechanisms is the knowledge of their origins and temporal ordering [1, 17-21]. Previous work utilized non-reversible genetic alterations in tumors, such as mutations and copy number changes, in either bulk or spatially resolved sequencing to track temporal events [22-26]. While these analyses are applicable to human tumor studies, they provide only inferences of chronological order or only monitor clonality and lack the precision to track associated cellular events.

Recent barcoding strategies in mammalian systems [27, 28], when combined with single-cell sequencing, have shown promise in unraveling the origins and chronological sequence of cellular events. However, their potential for recording temporal events long term is constrained by limited barcode diversities [29] and loss of information due to large deletion of multiple adjacent cut-sites [30, 31]. We are curious about the potential of a multimodal framework, pairing long term temporal tracking in mice with human single-cell multi-omics data, for addressing questions regarding cellular origins. For this, we developed a custom multi-purpose single-cell platform called **N**ative **s**gRNA **C**apture and **seq**uencing (NSC-seq) for simultaneous capturing of mRNAs and gRNAs, that can leverage self-mutating CRISPR barcodes (hgRNAs/stgRNAs) [32-34] to enable lineage tracking and temporal recording using accumulative mutation patterns. We utilized NSC-seq to decipher canonical developmental branching during mouse gastrulation. We demonstrated the ability of this platform to identify novel embryonic progenitor cell populations and new routes of cellular differentiation, as well as to provide fresh insights into the timing of tissue diversification. These results lay the foundation for *in vivo* multimodal recording for a wide variety of applications. We further leveraged this tracking approach by pairing it with genome-scale analysis of human tissues to illuminate the cellular origins of colorectal cancer. As part of the HTAN, we collected one of the largest multi-omic atlasing datasets on human sporadic polyps to date, comprising 116 polyps with scRNA-seq data and 418 polyps with mutational data. Paired analysis of human atlasing data, in conjunction with mouse intestinal tumor models, revealed the polyancestral origins of colorectal tumorigenesis. Our multimodal framework employing natural genetic changes in human paired with induced genetic changes in the mouse illuminate the complexities of cellular origins and temporal transitions, and their significance in early tumorigenesis.

### A temporal recording platfom

To enable CRISPR-based temporal recording at single-cell resolution, one must address the challenges of 1) non-polyadenylated hgRNA (but also sgRNA, stgRNA) capture, and 2) sparsity of single-cell data. We developed custom capture of non-polyadenylated gRNA that does not require redesign of whole gRNA libraries [35] or indirect readouts [36-38] (Fig. 1a and Extended Data Fig. 1a). Nearly 80% of gDNA mutations were detected in hgRNA with NSC-seq (Fig. 1b). We demonstrated hgRNA mutations were equivalent to gDNA ones for lineage tree reconstruction using controlled cell passage experiments (Extended Data Fig. 1b). Adaptation of NSC-seq to single-cell resolution using inDrops [39] demonstrated gRNA detection in 95% of identified single cells in cell line experiments, with their paired transcriptome data exhibiting similar quality as regular inDrops (Fig. 1c-d and Extended Data Fig. 1c-f, Supplemental methods). The effectiveness of using single-cell data is diminished by data sparsity, as illustrated by the applications to track cell division history and construct lineage trees. Previous work [5] and our results here show that gDNA barcode mutation frequency - as defined by the ratio of mutated vs. wild-type barcodes - tracks linearly with cell or organoid culture time when measured in bulk (Extended Data Fig. 2a-c). Due to single-cell data sparsity, only a fraction of barcodes can be detected on a per cell basis, invalidating the mutational frequency metric for single-cell use. We found that mutation density - as defined by the average number of mutations within barcodes - is immune to data sparsity and also tracks with time in organoid cultures and the renewing intestinal epithelium (Extended Data Fig. 2d-e). We revealed that mutational density increases at a faster rate in intestinal organoid cultures than the *in vivo* intestinal epithelium (Extended Data Fig. 2f), confirming that epithelial cells in organoid conditions are more proliferative [40]. Cellular turnover rates of common intestinal cell types, as inferred by mutational density, were also consistent with current knowledge (Extended Data Fig. 2g). Specifically, tuft cells exhibited a multimodal distribution of mutational densities, consistent with a heterogeneous cell population with different lifetimes [41, 42] (Extended Data Fig. 2h). NSC-seq applied to three murine embryonic time points to profile hgRNAs and mRNAs simultaneously also showed mutation density to increase over time (Fig. 1e-f), driven by cell type-specific changes (Extended Data Fig. 2i-j), but with no cell type bias in Cas9 expression or NHEJ activity (Extended Data Fig. 2k-l). While mutation density per barcode can be used for timing assessments, non-overlapping gRNA barcode expression detected per cell also limits information content used for cell phylogeny reconstruction. We thus augmented hgRNA mutational information with somatic mitochondrial variants (mtVars) [43, 44]. Briefly, we filtered out germline mtVars using a custom ‘germline mtVars bank’ (Supplemental methods), and then defined a lineage-determining cutoff from mtVar distributions using paired hgRNA mutations as ‘ground truth’ somatic variants (Extended Data Fig. 3a-d). Using this pipeline, we showed that mtVars also consistently increased over three embryonic time points (Fig. 1g-h), similar to hgRNA mutations (Fig. 1f). We further delineated the known developmental order of different murine brain layers prior to left/right brain segregation (Extended Data Fig. 3e) [45], and verified previously reported clonal relationships between 3 human breast cancer regions (Extended Data Fig. 3f), solely using mtVars on published spatial data [46, 47]. Single-cell analysis using hgRNA, mtVars, or both were able to accurately distinguish lymphoid and myeloid cells as distinct lineages in PBMCs (Extended Data Fig. 3g-j) and differentiate embryonic tissue types (Fig. 1i). Together, we developed a comprehensive pipeline of temporal and lineage tracking that is compatible with paired single-cell transcriptomic analysis (Fig. 1j).

**Fig. 1:**
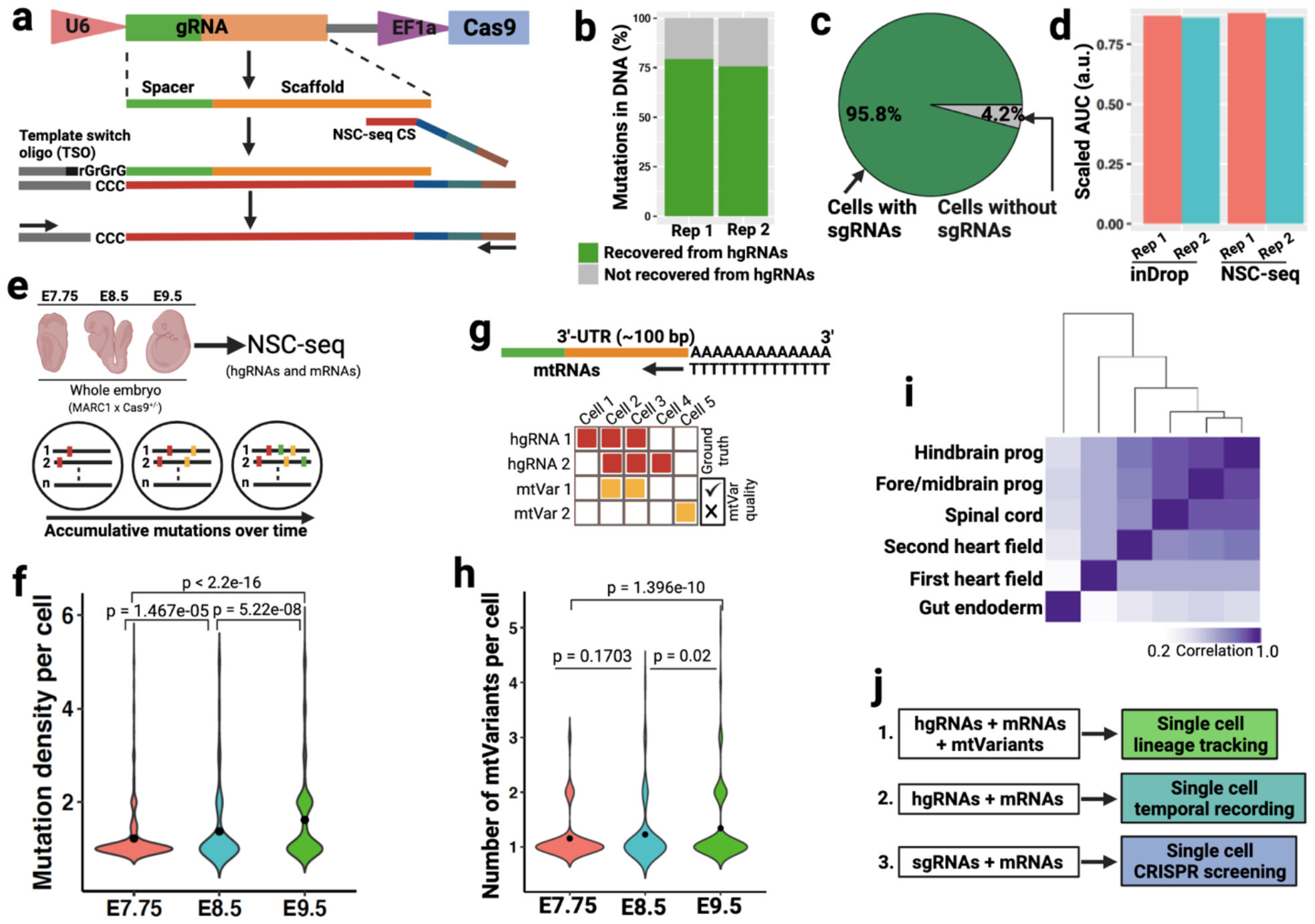
Optimization of a multi-purpose single-cell capture platform. (**a**) Guide RNA (gRNA) capture schematic for the NSC-seq platform. Target site of gRNA scaffold anneals to NSC-seq capture sequence (CS) with a cellular barcode (blue) and UMI (green). An additional sequence (gray) is added to the 3’-end of the cDNA via template switching during reverse transcription to enable downstream library amplification. This gRNA capture approach is compatible with any type of gRNAs (sgRNA, hgRNA, and stgRNA) that contains the target site sequence in the scaffold (see Extended Data Fig. 1). (**b**) Cas9-induced mutations recovery by direct hgRNA capture as compared to mutations detected in DNA of the same samples. (**c**) Guide RNA capture efficiency by NSC-seq assessed in an experiment where all cells from a drug-selected cell line should contain sgRNAs. (**d**) Comparative transcriptome capture efficiency between standard inDrops [39] and NSC-seq experiments. (**e**) NSC-seq experiments on developmentally barcoded whole embryos where Cas9 is constitutively expressed (top). Accumulative mutations on homing barcode regions increase over time (bottom) [5, 32]. (**f**) Average mutation density over embryonic time points (see Extended Data Fig. 2a). Black dot represents geometric mean for each time point and p-value derived from Student’s t-test. (**g**) Somatic mitochondrial variants (mtVars) calling from mitochondrial RNA (mtRNA) (top) [43, 44]. Approach to filter informative mtVars for lineage tracking using hgRNA mutations as ‘ground truth’ (bottom) (see Extended Data Fig. 3b-d). (**h**) The Number of somatic mtVars per cell over embryonic timepoints. Black dot represents geometric mean for each timepoint and p-value derived from Student’s t-test. (**i**) Pearson correlation coefficient heat map of variant proportions combining hgRNAs and mtVars for selected tissue types presented as pseudobulk from an E9.5 embryo (see Extended Data Fig. 4). (**j**) Multi-modal application of the NSC-seq platform.

**Fig. 2:**
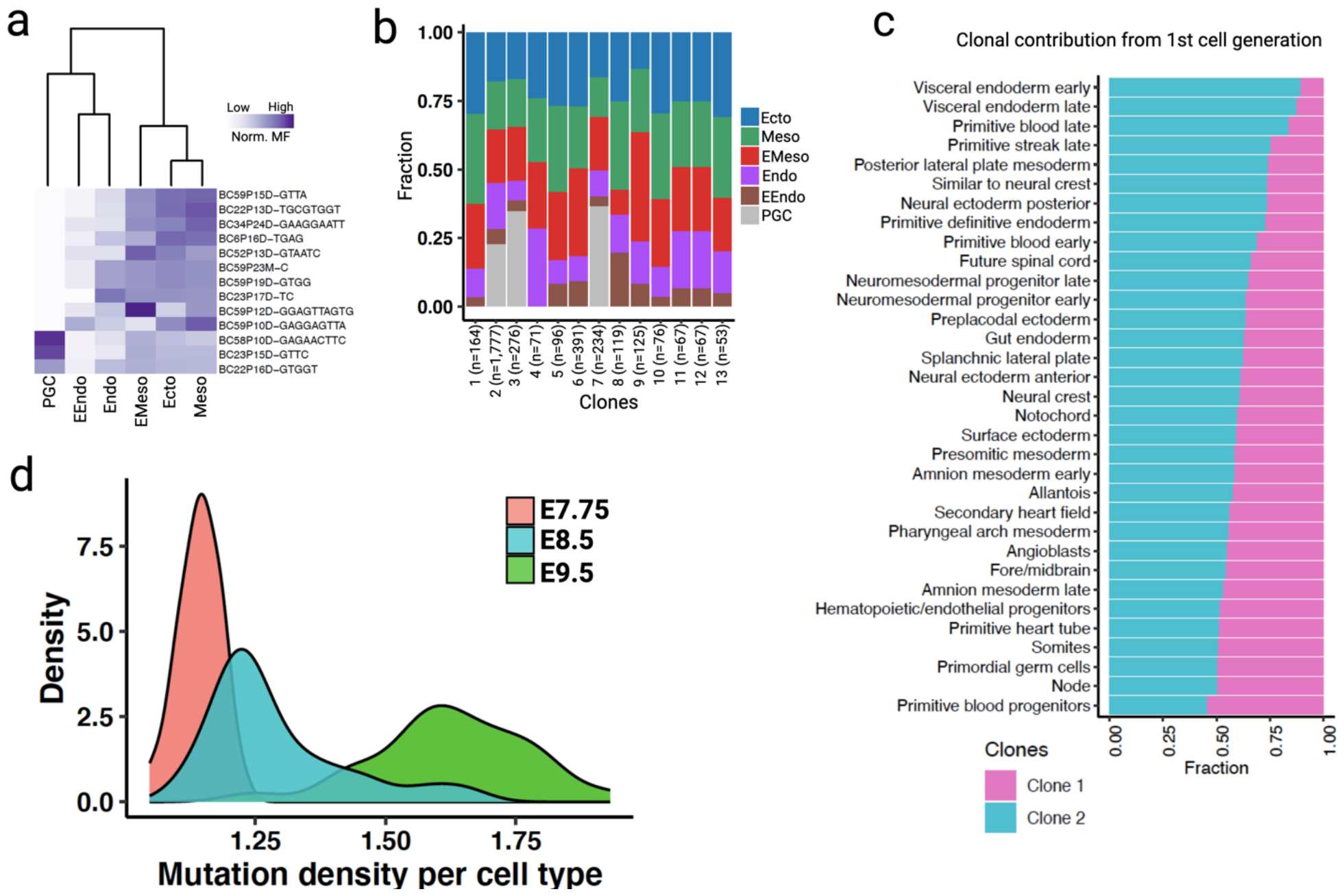
Lineage and temporal recording of mouse embryogenesis. (**a**) Normalized mosaic fraction (MF) of early embryonic mutations (EEMs) heat map for E7.75 embryo to reconstruct lineage relationships within the major germ layers (see Extended Data Fig. 5 and 7). (**b**) Contribution of different EEMs towards various germ layers at E7.75. (**c**) Clonal contribution from a 1^st^ cell-generation mutation (Clone 1) at E7.75 across individual tissue types (p = 1.57e-13, Kolmogorov-Smirnov test for the null hypothesis of symmetry) compared to all other clones aggregated as “Clone 2” (see Extended Data Fig. 5l-m). (**d**) Density plots representing cumulative turnover of different tissue types across three embryonic timepoints. The widths of the mutation density distributions represent the variation by which different cell types have proliferated across timepoints (see Extended Data Fig. 6 and Supplemental table 2 for mutation density per cell type). Ecto, embryonic ectoderm; Meso, embryonic mesoderm; Endo, embryonic endoderm; EMeso, extra-embryonic mesoderm; EEndo, extra-embryonic endoderm; PGC, primordial germ cell.

**Fig. 3:**
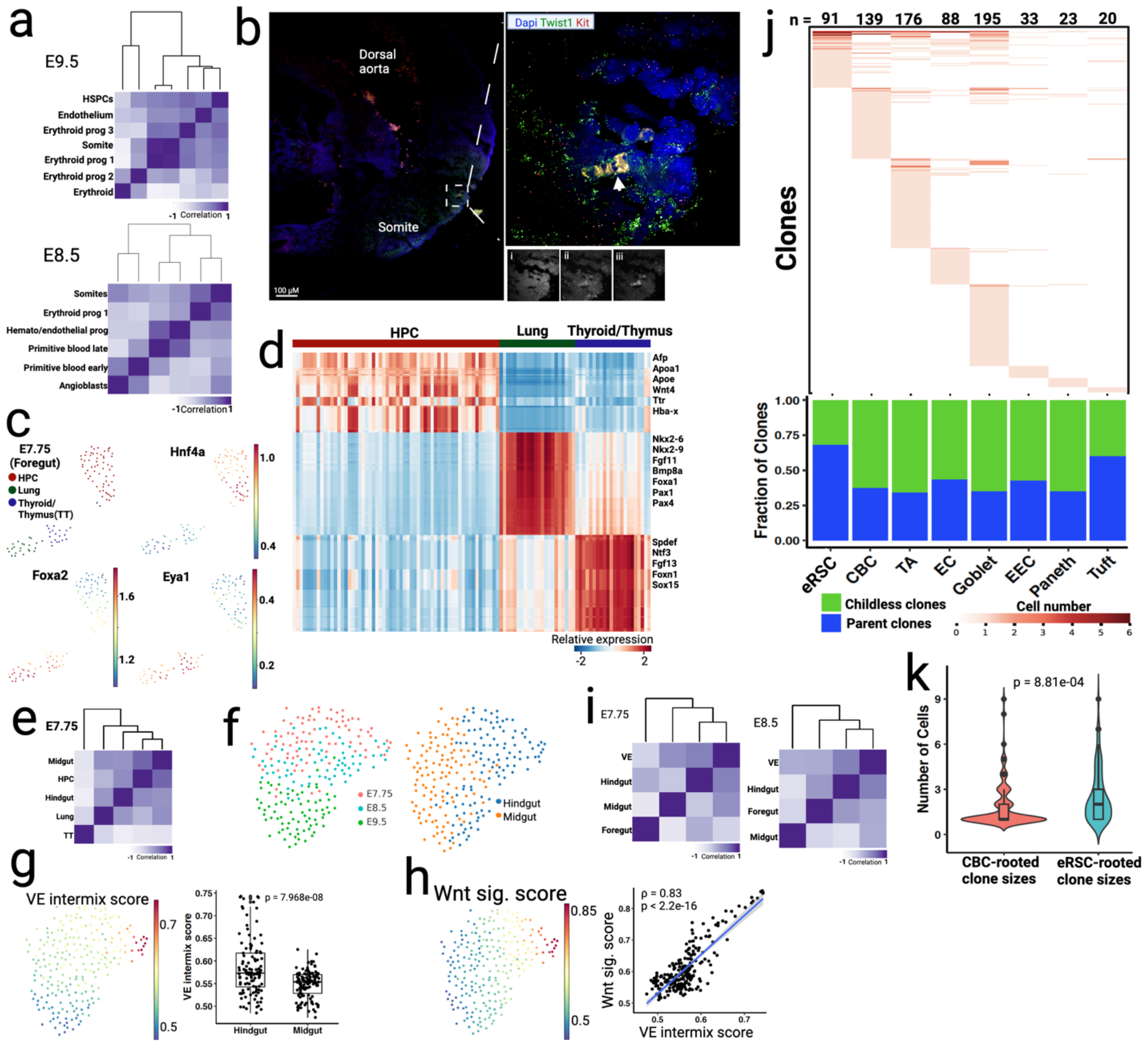
Embryonic lineage diversification and gut development. (**a**) Pearson correlation coefficient heat maps of variant proportions (hgRNAs and mtVars) presented as pseudobulk within hematopoietic and somite cell types from E9.5 (top) and E8.5 (bottom) embryos. (**b**) Multiplex HCR-FISH staining of somite (Twist1) and hematopoietic (Kit) markers in a E9.5 embryo. A cluster of hematopoietic cells (white arrowhead) in somite area is shown in inset image (right). Inset mono-color images represent Dapi (i), Twist1 (ii), and Kit (iii) (see Extended Data Fig. 8). (**c**) Force-directed layout of foregut cells (E7.75) that are colored by annotated tissue types. HPC, hepatopancreatic cells. Gene expression of HPC (Hnf4a), lung (Foxa2), and thyroid/thymus (Eya1) markers are overlaid here. (**d**) Heat map of differentially expressed genes among three foregut tissue types at E7.75. Tissue type-specific genes are labeled on the right. (**e**) Pearson correlation coefficient heat map of distinct tissue types from gut region (E7.75) presented as pseudobulk, similar to panel a (see Extended Data Fig. 9). (**f**) Force-directed layout of midgut and hindgut cells are colored by embryonic time points and regions. (**g**) Visceral endoderm (VE)-intermix score overlay onto f. Quantification of the VE intermix score in hindgut compared to midgut cells (Student’s t-test). (**h**) WNT signaling score overlaid onto f. Correlation analysis between the WNT signaling score (y-axis) and VE-intermix score (x-axis). (**i**) Pearson correlation coefficient heat maps of gut regions with VE as pseudobulk from E7.75 and E8.5 embryos. See Extended Data Fig. 10c for VE annotation. (**j**) Distribution of clones across cell types in adult mouse small intestinal epithelium (see Extended Data Fig. 11). Number at the top represents the total number of detected clones per cell type. Heat map color represents the number of cells found comprising a clone within a given cell type. A plot (below) showing the fraction of parent and childless clone comprising each cell type, as defined in Extended Data Fig. 10j. EEC, enteroendocrine cells; CBC, crypt-based columnar cells; eRSC, embryonic revival stem cells; EC; enterocytes; TA, transit-amplifying cells. (**k**) Violin plots of CBC-rooted and eRSC-rooted clone sizes.

### Embryonic lineage and cell division tracking

We then more deeply analyzed combined single-cell barcoding and transcriptome data of E7.75, E8.5, and E9.5 embryos to glean biological insights from NSC-seq. Cell type annotation using conventional gene expression analysis revealed canonical cell types and germ layers at each of the time points [27, 48, 49] (Extended Data Fig. 4, Supplemental information). Notably, more defined cell types emerged at E9.5 compared to earlier time points (E7.75/8.5), consistent with the established timeline of mammalian development. This distinction prompted two separate sets of cellular annotations (Extended Data Fig. 4a-h). Matching our data with previously generated scRNA-seq data at E7.0 and E8.0 supported the correct development timing of our single-cell embryonic data (Extended Data Fig. 4i), and data quality was typical of this experimental platform (Extended Data Fig. 4j-l, Supplemental methods). Regarding the quality of barcode mutations called, the distribution of mutations amongst cells, the frequency of different types of mutations, the incidence of random collision mutations, the number of mutations as a function of cell type, and the barcode lengths were all consistent with previous reports (Extended Data Fig. 5a-e) [21]. Early embryonic mutations (EEMs) occur during the initial stages of cell division in development and are inherited by a significant portion of cells within the embryo (Extended Data Fig. 5f-g). The proportional presence of these mutations amongst cells, referred to as the mosaic fraction (MF), can serve as an indicator of the cell generation (CG) when these mutations originated (Extended Data Fig. 5h-i) [50]. Progressive restriction of EEMs shared in tissues enable the use of MFs to model early divergence of germ layers and tissue types (Fig. 2a). Mouse primordial germ cell (PGC) lineage segregated from other embryonic and extra-embryonic lineages, supporting possible early allocation of cells to the PGC lineage that has also been reported in mouse [51] and human [52, 53]. We also found a similar MF between mesoderm and ectoderm that supported a shared progenitor population, as reported before [54]. Notably, extra-embryonic endoderm (EEndo) and embryonic endoderm (Endo) appeared to share origins, although they are known to originate from two distinct tissue layers, hypoblast and epiblast, respectively. However, there is literature supporting some degree of shared progenitors, lineage convergence, and intermixing between these tissues [27, 48, 55-58]. We also assessed the clonal contributions of different EEMs towards germ layers (early) or tissue types (late) and observed unequal contribution between different early clones (Fig. 2b and Extended Data Fig. 5j-k). We found unequal partitioning of first cell generation clones across different tissue types (Fig. 2c, p = 1.057e-13), suggesting that the specific lineage commitment of early embryonic progenitors is not predetermined, but rather subject to potential induction or stochastic processes (Extended Data Fig. 5l-m). This phenomenon has been previously reported in mammals [7, 8, 25, 53, 54], and is not observed in *C. elegans*.

**Fig. 4:**
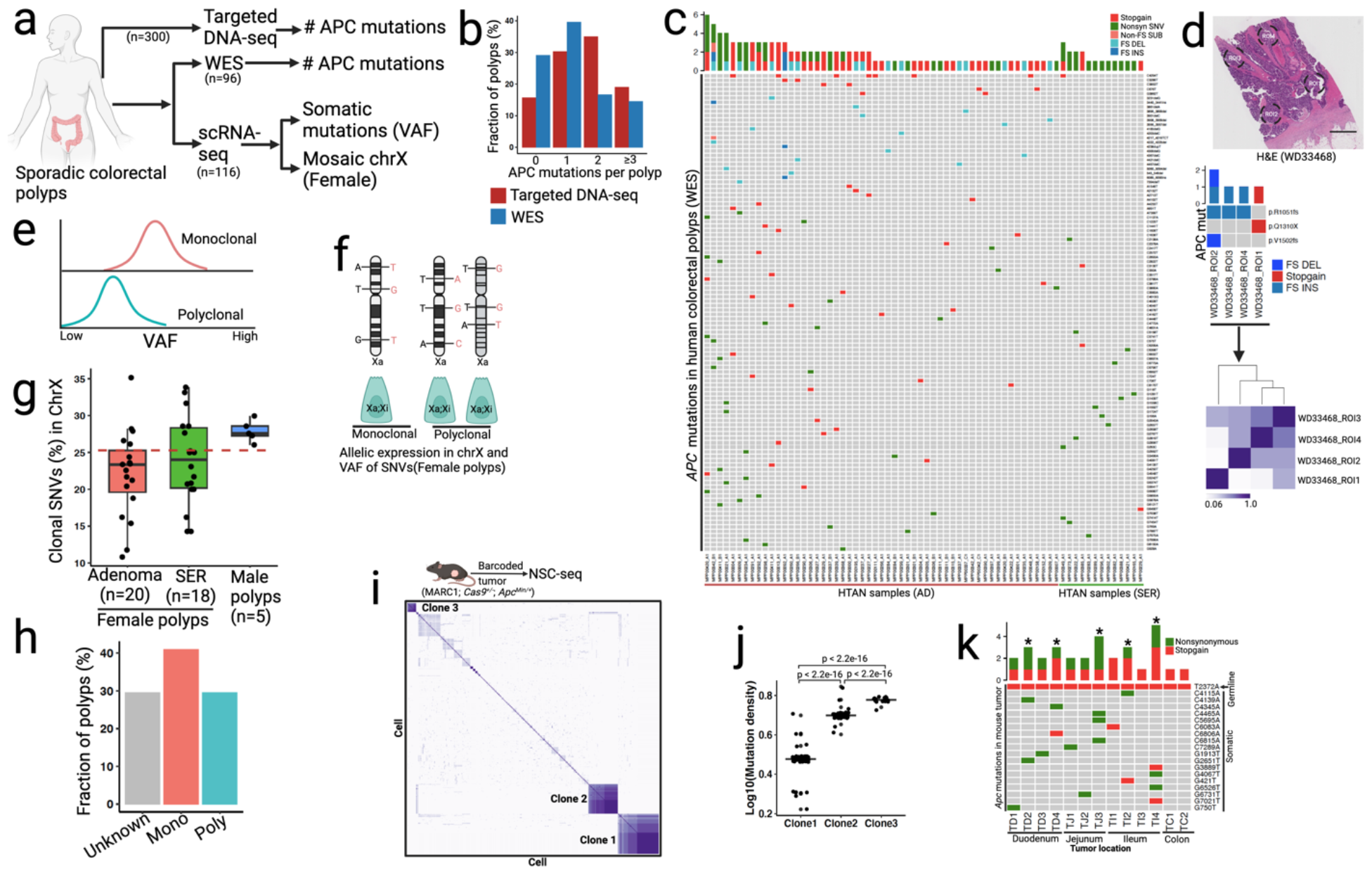
Clonal origin of colorectal precancer. (**a**) Overview of experimental design for profiling clonal origin across multiple human datasets. (**b**) Bar plots summarize the number of APC mutation per polyp using targeted DNA sequencing and whole-exome sequencing (WES). (**c**) OncoPrint plot represents the number of APC mutations across human polyps using WES. Here we only show polyps with at least one deactivating APC mutation. See Extended Data Fig. 12 for more details. (**d**) Multi-region (punch biopsy) WES of a human CRC sample represents distinct APC mutations and Pearson correlation coefficient heat map (bottom) of somatic mutations within the regions of interest (ROI) [23]. (**e**) Expected median VAF distribution under different clonal architectures. (**f**) Mosaic X chromosome inactivation patterns in female polyps can delineate the clonal origin of cells using expression-based X-linked somatic clonal SNVs. Male polyps are considered as monoancestrol due to single X chromosome in male. See Extended Data Fig. 12f-h and supplemental method section. (**g**) Box plots represent distribution of X-linked clonal SNVs (%) between male and female polyps. Red line is a cut-off to assign ancestry in female polyps. See Extended Data Fig. 12i-j. (**h**) Summary of median VAF-based polyp profiling. (**i**) Pearson correlation coefficient heat maps of variants (hgRNAs and mtVars) from mouse intestinal tumor (*Apc*^*Min/+*^)-derived single cells. Distinctly correlated regions are marked by three clones within the same tumor. See Extended Data Fig. 13. (**j**) Estimated mutation density for the three assigned clones in i. (**k**) OncoPrint plot represents the number of *Apc* mutations across mouse tumors using WES. Five tumors (asterisk) show more than 2 distinct *Apc* mutations.

The regulation of organ size is one of the most fundamental processes of embryonic development, primarily governed by organ-specific cell division rates, and to a lesser extent, rates of apoptosis [59-62]. Here, we developed a catalog of cell division histories of different organs to reveal insights into the timing and scale of cell division across tissues during development (Supplemental methods). Using mutations within NSC-seq barcodes, we quantified the cumulative number of cell divisions per tissue type at three gastrulation time points (Extended Data Figure 6a-b, Supplemental Table 2). We observed that the relationship between the number of cell divisions and the known tissue mass differs among various tissue types, which could be attributed to a number of variables, including differential progenitor field size, the timing of progenitor specification, cell death, cellular lifespan, and cell competition across tissue types [63]. Additionally, our data revealed a widening distribution of tissue-specific cumulative cell division at both E8.5 and E9.5 stages whereas a narrow unimodal distribution was observed for the E7.75 stage (Figure 2d), suggesting that tissue-specific cell division and diversification initiates after the E7.75 stage. In general, we observed high proliferation of hematopoietic progenitors during gastrulation, while cardiomyocytes and endothelium showed low proliferation (Extended Data Fig. 6a-b). We noticed an emergence of various intermediate hematopoietic progenitors at E9.5 with distinct cellular turnover histories, supporting diverse roots of hematopoiesis during early embryonic development as previously reported [64-70]. Cumulative cell division levels for forebrain progenitors were higher than hindbrain progenitors (Extended Data Fig. 6b), supporting known turnover kinetics to maintain the relative size of brain regions during mammalian neurogenesis [63, 71-73]. In addition, we found a constant rate of cell proliferation for gut endoderm over embryonic time points, a rate similar to the turnover of the adult intestinal epithelium (Extended Data Fig. 6c and 2e). Overall, differential proliferation timing and kinetics between organs during gastrulation were observed. These variations occasionally corresponded to the organ size, but not always. We also demonstrated that for certain tissues, proliferation rates were set during gastrulation and persisted throughout life [53]. Overall, this catalog serves as a basis for studying embryonic cellular proliferation kinetics and adds a temporal axis in lineage diversification [1] to complement lineage tracking.

Next, a single-cell phylogenetic reconstruction (Supplemental methods) was conducted using NSC-seq data at a higher information content per cell than previous approaches (Extended Data Fig. 7a-c). Pseudo-bulk reconstruction of embryonic tissue relationships generally recapitulated canonical knowledge of germ layer development (Extended Data Fig. 7d). Phylogenetic distance analysis from single cell tree supports the closer proximity of EEndo to root compared to Endo or Meso to root (Extended Data Fig. 7e). A wider distribution of the phylogenetic distances across cell type was observed at E8.5 and E9.5 compared to E7.75 (Extended Data Fig. 7f), supporting the initiation of tissue-type diversification after E7.75 illustrated above (Fig. 2d) [74]. Furthermore, computational inference from single-cell lineage tree topology (Supplemental methods) estimated the number of epiblast progenitors (n=∼28) and extrapolated unequal progenitor field size between ectoderm and mesoderm stemming from these progenitors (Extended Data Fig. 7g-h) [75].

### Instances of unconventional embryonic lineage diversification

We highlight three examples of unconventional lineage diversification we observed during embryonic development. Lineage analysis at both E8.5 and E9.5 indicated that Erythroid Progenitor 1 (EryPro1) share common ancestry with somite (Fig. 3a). We then reanalyzed somite, endothelium, and hematopoietic cell types, all potential progenitors to EryPro1, and found that EryPro1 did not express yolk sac (Icam2, Krd, and Gpr182), endothelial (Pecam1), and embryonic multipotent progenitor (eMMP) markers (Flt3) (Extended Data Fig. 8a-c) [67]. In contrast, EryPro1 expressed somite-specific markers (Twist1, and Sox11) and showed upregulation of WNT signaling, which comprised an EryPro1-specific gene signature (Extended Data Fig. 8d-f, Supplemental table 3). Additionally, RNA velocity, MF of EEMs, and clonal analyses all supported a developmental relationship from somite to EryPro1 (Extended Data Fig. 8g-i). Indeed, multiplex HCR-FISH of somite and erythroid markers revealed a cluster of Kit+ erythroid cells in the somite region of the E9.5 embryo (Fig. 3b), supporting a somite-derived erythroid progenitor population. The EryPro1 population is present at E8.5, but not at E7.75 stage, whereas somite cells were observable at E7.75 (Extended Data Fig. 8j-m). Gene expression analysis showed some somite cells from E8.5 co-expressed hematopoietic transcription factors (Gata1 and Gata2) and low level of hemoglobin gene (Hbb-bt), implying a cell-state transition from somite to EryPro1 (Extended Data Fig. 8n-o) [76, 77]. Finally, pseudotime analysis revealed a distinct developmental trajectory from somite to EryPro1, in addition to the expected trajectory from somite to sclerotome (Extended Data Fig. 8p). Thus, our data supports a somite-derived hematopoietic population during late gastrulation of mammalian development, similar to zebrafish [66].

We next sought to understand gut endoderm development in context of regionalization and progenitor specification timing. Endoderm (definitive and visceral) cell populations from E7.75 and E8.5 embryos were plotted together to reveal region-specific markers as early as E7.75, implying regionalization (spatial patterning) at that early time point (Extended Data Fig. 9a-d). We then focused our analysis on region-specific progenitors of the gut at E7.75. Analysis of foregut population from E7.75 revealed three distinct clusters: hepatopancreatic (HPC) progenitors (Hnf4a+), lung progenitors (Foxa2+), and thyroid/thymus (TT) progenitors (Eya1+) (Fig. 3c). Gene expression, regulon activity, and lineage analysis showed that the HPC population is relatively distinct from lung and TT progenitors (Fig. 3d-e, Extended Data Fig. 9e-f). Similar progenitor populations from the foregut were found at E8.5 (Extended Data Fig.9g-h) but not at E7.5 (Extended Data Fig.9i), implying precise timing of progenitor specification at E7.75. Analysis of the remaining definitive endoderm populations similarly revealed distinct gene expression between midgut (Gata4, Pyy, and Hoxb1) and hindgut (Cdx2, Cdx4, and Hoxc9) progenitors as early as E7.75 (Fig. 3f, Extended Data Fig.9j). Regulon analysis also suggested distinct region-specific activities for midgut (Gata4, Foxa1, and Sox11) and hindgut (Cdx2, Sox9, and Pax2) progenitors at this time point (Extended Data Fig. 9k). Pseudo-time and CytoTRACE analyses resulted in a logical developmental trajectory from E7.75 to E9.5 (Extended Data Fig. 9l). We found notable region-specific differences in WNT and BMP signaling over developmental pseudo-time (Extended Data Fig. 9m). Significantly higher WNT signaling activity was observed in hindgut compared to midgut progenitors at E7.75 (Extended Data Fig.9n-o). Consistent with the literature, the WNT target gene Lgr5, a canonical intestinal stem cell marker [78], was highly expressed in hindgut [79], whereas Lgr4 and Lgr6 were expressed in the midgut (Extended Data Fig. 9p). Our results revealed early differential usage of developmental signaling pathways between progenitors of different regions, supporting an early progenitor specification model during endoderm development [80, 81].

We also examined the lineage relationship between visceral endoderm (VE) and definitive endoderm (DE) during embryonic development. We derived a VE score using reported VE infiltration-specific marker genes and showed that the score could accurately mark sorted VE-derived cells (Extended Data Fig. 10a). Application of this score to our data identified cells that demonstrated high VE-intermixing in the developing hindgut (Fig. 3g, Extended Data Fig. 10b). Unexpectedly, we found that the VE-intermixing score correlated with WNT signaling score and genes (*Lgr5, Axin2*, and *Fzd10*) (Fig. 3h, Extended Data Fig. 10c), which was supported by higher *Lgr5* expression in sorted VE-derived cells than DE-derived cells (Extended Data Fig. 10d). Multiplex HCR-FISH showed the presence of cells with co-expressing VE-marker gene *Cthrc1* and *Lgr5* in the posterior gut region (dotted line) (Extended Data Fig. 10e). Lineage analysis using mutational barcodes supported a lineage relationship between hindgut and VE, likely resulting from VE-derived cells mixing into the hindgut during gastrulation (Fig. 3i). This relationship persists at E9.5, as supported by differential lineages between midgut and hindgut (Extended Data Fig. 10f-g). To determine the role of VE-derived cells in post-gastrulation, we analyzed midgut and hindgut tissues at the E14.5 time point and found that hindgut epithelium has a higher VE intermix score than the midgut epithelium (Extended Data Fig. 10h-i), consistent with the results above. We then assessed the ability of these cells to contribute to epithelial development by performing a ‘parent-childless’ clonal analysis using a reported approach [17] (Extended Data Fig. 10j). VE-derived cells have a high parent clone fraction, implying that they have a higher potential to give rise to progeny (Extended Data Fig. 10k). Mutation density analysis also demonstrated VE-derived cells have accumulated more divisions at E14.5 compared to other DE-derived cells, demonstrating their post-gastrulation activities (Extended Data Fig. 10l). Finally, we performed mutational barcode analysis of adult tissues derived from foregut, midgut, and hindgut and found that hindgut-derived tissues maintain a separate lineage branch from midgut- and foregut-derived tissues even into adulthood (Extended Data Fig. 10m). Thus, our data support previous reports of VE-derived cells intermixing with DE (Extended Data Fig.10n) predominantly in hindgut [48], and their potential contribution to gut epithelial development [27, 48, 55-58].

### Identification of an early embryonic progenitor of gut epithelium

NSC-seq applied to the adult gut identified a unique cell population related to enterocytes in the small intestine, which we labeled embryonic revival stem cells (eRSCs) (Extended Data Fig. 11a-c). A gene signature derived from this cell population was also able to identify the same cells in another publicly available dataset (Extended Data Fig. 11d-e). Mutational lineage analysis demonstrates a developmental relationship between crypt-based columnar cells (CBCs) and eRSC cells, indicating they potentially derive from each other (Extended Data Fig. 11f). However, eRSCs exhibit a significantly higher mosaic fraction, implying that they are derived from much earlier cell generations compared to CBCs, which develop relatively late during fetal intestinal development [82] (Extended Data Fig. 11g). A smaller number of progenitors that give rise to these cells inferred from single-cell lineage tree topology (Extended Data Fig. 11h) supports their earlier specification stemming from fewer progenitors available at earlier development. Clonal contribution analysis using hgRNA mutations demonstrates that the eRSCs population possesses a larger clone size, thus contributing more progenies to the intestinal epithelium than CBCs (Fig. 3j-k). This result was repeatable, supporting that the eRSC population acts as a stem/progenitor-like population during intestinal development (Extended Data Fig. 11i-n). We selected Transducer of Erbb2.2 (Tob2) as a marker of eRSC cells and found that they are located at the bottom of the adult small intestinal crypt via immunofluorescence analysis (Extended Data Fig. 11o-p). These data support that eRSCs can act as stem/progenitor-like cells to populate the gut during embryogenesis, in contrast to the limited contribution of the CBC population at that time [82].

### Clonal analysis of colorectal precancers

We next tackled the question of clonality during tumor initiation in the gut. While individual crypts comprised of a single clone, different ancestral models of tumorigenesis have been proposed and are still being debated [83]. The prevailing model, with support from human colorectal cancer data, is the monoclonal model, where a tumor is initiated from a single stem cell residing in a crypt [84, 85]. However, selection and clonal sweeps that occurred in advanced cancers tend to erase clonal histories occurring earlier in tumorigenesis [86]. Further, lineage tracing studies in the mouse have shown that some tumors can be initiated from multiple ancestors, resulting in tumors with multiple lineage labels [87, 88]. An accompanying paper in this issue leveraging intra-patient embryonic clone sharing amongst multiple familial adenomatous polyps (FAPs) within the same patient demonstrates the possibility of polyancestry in tumor formation [89]. While embryonic clone mixing can only be leveraged in hereditary diseases such as FAP, we sought to find evidence of polyancestry in sporadic human polyps. We expect polyancestry to only occur in a minor subset of polyps, thus requiring a large sample size analysis for our study. Thus, we collected new scRNA-seq data, resulting in a total of 116 polyp datasets (AD=70, SER=42, UNK=4) from 3 different cohorts of patients at Vanderbilt University Medical Center (VUMC) [90] (Fig. 4a, Extended Data Fig. 12a). Out of these, 96 polyps (AD=63, SER=33) had matching WES data. These data were generated from distinct regions of the colon from a distribution of 96 patients with diverse racial backgrounds and ages (Supplemental table 4). In addition, we analyzed targeted DNA sequencing from ∼300 polyps from Tennessee Colorectal Polyp Study (TCPS) to assess APC mutations [90]. Here, we present several analyses drawn from this human data to support conclusions regarding the clonality of colorectal tumor initiation.

*APC* is considered the gatekeeper gene in FAP and the majority of sporadic CRCs. Loss of function of both *APC* alleles, resulting in WNT pathway activation, is thought to initiate tumorigenesis [91]. Thus, the number of unique APC mutations can be used to assess clonality in conventional adenomas (ADs) [92]. In a diploid genome, a monoancestral adenoma should present at most two unique APC mutations that lead to loss of function of both alleles [93], given that there is no selective advantage for additional mutations. Using our TCPS data, we found that ∼20% of polyps show ≥3 unique APC mutations, implying more than one founder clone in those polyps (Fig. 4b, Extended Data Fig. 12b). Similar to these results, WES data from our VUMC polyp dataset showed potential polyancestry to occur at ∼15% of the polyps (Fig. 4b-c, Supplemental table 4). APC mutation analysis using multi-regional WES in a cohort of 23 colorectal carcinoma (CRC) samples from VUMC [23] showed only 1 specimen to exhibit potential polyancestry (Fig. 4d). This is consistent with the occurrence of clonal sweeps during tumor progression, as seen in external cohort datasets, that erases the clonal history of tumor initiation (Extended Data Fig. 12c) [94, 95].

To provide additional ancestry evidence, we called somatic SNVs from single-cell transcriptomics data of colorectal polyps using two independent pipelines (Extended Data Fig. 12d) [96, 97]. Clonal composition was then assessed using the variant allele frequency (VAF) distribution of somatic SNVs (Supplemental method). If a polyp is derived from a single founder clone, the VAF distribution of its somatic SNVs would be higher than that of a polyancestral polyp due to a higher fraction of shared SNVs across a single founder-derived population (Fig. 4e) [98-101]. We calculated median VAF from polyps (n=86) and found a wide variation across polyps, implying existence of both monoancestral and polyancestral polyps (Extended Data Fig. 12e). To establish a polyancestry cut-off based on VAF distribution, we leveraged the concept of X-linked inactivation in female polyps (n=46). During early embryonic development in females, one X chromosome in somatic cells becomes randomly silenced to balance X-linked gene dosage [102]. This pattern persists in daughter cells, creating a mosaic of inactivated X chromosomes in adult female tissues. Therefore, somatic SNVs within X-linked transcripts can be used as developmental markers to track the clonal origin of cells in females (Fig. 4f, Supplemental method) [103, 104]. In males with a single X chromosome, mosaic expression of X-linked genes is absent, and thus male polyps can stand in as “monoancestral” when solely considering X-linked SNVs (Extended Data Fig. 12f). We thus used simulations, mixing male polyps to establish baseline distributions of X-linked SNVs that distinguish between monoancestral and polyancestral polyps. As anticipated, the proportion of X-linked clonal SNVs decreased in relation to the degree of polyancestry (as simulated by the number of mixed male polyps) (Extended Data Fig. 12g). Examining female polyps on the same scale revealed a significant number of female polyps to potentially be polyancestral (Fig. 4g); many of those were also classified as polyancestral from APC mutation assessment (Extended Data Fig. 12h). A wide distribution of clonal X-linked SNVs in female polyps also indicated the potential for different numbers of founder clones (Fig. 4g). To extend analysis to all single-cell SNVs in addition to X-linked SNVs, we examined VAF distributions in female polyps that have been assigned as monoancestral or polyancestral based on X-linked SNVs. Assigned monoancestral polyps exhibited higher median VAF compared to polyancestral polyps, and we were able to establish a median VAF distribution cut-off of 0.20 to identify polyancestry (Extended Data Fig. 12i-j, Supplemental table 4). Applying VAF distribution analysis to all polyps, we found ∼29% of polyps to be polyancestral (Fig. 4h, Supplemental table 4), comparable to APC mutation-based assessment (Fig. 4b). Thus, analysis of multiple data types supports that a substantial subset of human colorectal precancers arose from multiple non-cancer ancestors.

For additional orthogonal confirmation, we applied WES data to a linear model that distinguishes between neutral and selective evolution (Extended Data Fig. 12k-l) [86]. We found that higher proportion of the assigned monoancestral polyps showed a signature of clonal selection (R^2^<0.98) compared to the assigned polyancestral polyps (Extended Data Fig. 12m). With some polyancestral tumors transitioning to clonal selection, about 60% of the polyps overall showed clonal selection by this analysis (Extended Data Fig. 12n), consistent with previous reports of selective pressures exerted during malignant progression [86, 94]. Moreover, adenoma-specific cells (ASC) of assigned monoancestral polyps showed higher expression of genes associated with cell cycle, nucleic acid synthesis, and protein translation signatures than polyancestral polyps, which can be attributed to a highly proliferative, stem cell-expansion phenotype that may drive clonal selection (Extended Data Fig. 12o-s) [105]. These data suggest that clonal selection can occur during the premalignant stage, and that increased selective pressures resulting in decreased clonality may be a hallmark of precancer to cancer transition.

Whereas data analysis can infer ancestry from human tumors, our single-cell barcoding mouse model provides a valuable resource for validating clonal composition of tumors *in vivo*. We generated barcoded tumors from the *Apc*^*Min/+*^ murine model of colorectal cancer where tumorigenesis occurs as a result of random mutations inactivating the second allele of *Apc*. We found that the tumor is comprised of both normal and tumor-specific cells, similar to human adenomas (Extended Data Fig.13a-c, Supplemental methods) [90]. Evaluating tumor-specific cells using NSC-seq demonstrated an increased proliferation signature, stemness, fetal gene expression (*Marcksl1*), and clonal contribution compared to normal CBC stem cells (Extended Data Fig.13d-e), consistent with the tumorigenicity of these cells. Examination of phenotypically normal cells within the tumor showed normal-like progenies of tumor-specific cells which can be distinguished from their normal counterparts by their higher barcode mutation densities and shared barcode mutation profiles with tumor cells (Extended Data Fig. 13f). Interestingly, these progenies consisted of enterocytes and Paneth cells, consistent with WNT-restricted aberrant differentiation of intestinal tumor cells [106]. To delineate clonality, we first used shared barcode mutations in lymphocytes, demonstrating that tumor infiltrating lymphocytes have expanded clonally compared to peripheral blood lymphocytes, which were mostly polyclonal (Extended Data Fig.13g). Similar analysis revealed three founder clones within tumor-specific cells (Fig. 4h). The three clones were distinct in many characteristics, including mutation density, clonal contribution, biased differentiation, and various gene expression signatures (Fig. 4i, Extended Data Fig. 13h-k). More importantly, single-cell phylogenetic analysis showed independent tumor founder clones to arise from distinct normal epithelial ancestors (Extended Data Fig. 13l). We also performed WES and found 5 of the 13 murine intestinal tumors have ≥3 unique mutations in the *Apc* gene, implying multiple founder clones similar to human adenomas (Figure 4j). Moreover, ∼ 40% of murine tumors showed evolutionary selection pressure comparable to human adenomas (Extended Data Fig. 13m). The normal cell-of-origin of tumor cells can also be examined by early embryonic clonal intermixing using barcode mutations in both tumor and adjacent normal tissues from the same mouse [107-110]. Early embryonic clonal intermixing was seen in 4 out of 5 mouse polyancestral tumors (Extended Data Fig. 13n-o, Supplemental table 4). Thus, our data showed instances of widespread EEM sharing, indicating that barcode mutations used to determine polyclonality were also found in adjacent normal cells. Taken together, our results generated from human sporadic polyps and validated in a mouse model provide insights into the evolutionary dynamics at the earliest stage of tumorigenesis in the mammalian colon.

## Discussion

Identifying origins of cells is an important endeavor in both developmental biology and cancer studies. This challenge becomes particularly pronounced when the progenitor cell is manifested within a specific subset of a given cell type. As an example, tumors can arise from a subset of normal cells in a seemingly random fashion [111] or under the influence of factors that push them towards this fate [112, 113]. Using single-cell genomic information from 116 human colorectal polyps, we present orthogonal evidence from different analyses to demonstrate the substantial number of instances where colorectal polyps emerge from multiple distinct clonal origins. Results from this study and the companion study by Schenck et al. [89] support wide prevalence of polyancestral composition of human polyps in both genetic and sporadic settings. This finding in the gut is in line with recent reports on polyancestral human breast cancer initiation [101, 114].

Considering that advanced cancer typically presents as monoclonal, and clonal selection can be observed in some, but not all, polyps, it raises an intriguing possibility that the subset of polyps undergoing a selection process may be primed to progress to advanced cancer. Hence, future research may elucidate whether clonality can serve as a predictive biomarker for precancers that will advance to malignancy, in contrast to polyps that maintain polyclonality. Nevertheless, approaches to functionally study the origins of predetermined cell fates in model systems are lacking. Here, we additionally leveraged clonal progeny generated by synthetic barcode mutations in a single-cell platform (NSC-seq) to enable retracing cell lineage origins backwards in time.

We first applied this lineage tracking platform to study mammalian development over different time scales from zygote to adult. Our analysis of gut endoderm development revealed regionalization of endoderm and progenitor specification to initiate earlier than previously appreciated, and suggested that these two processes may occur simultaneously [80]. In addition, our gut lineage analysis showed convergence of cells from extra-embryonic origin to an embryonic endoderm state, supporting previous observations [27, 48, 55-58], and extended the contribution of extra-embryonic cells to gut epithelial development. Moreover, temporal analysis of embryonic development revealed a significant shift in tissue-specific cell expansion after E7.75. Hence, our study provides clues about developmental timing of lineage diversification that can prompt studies into extrinsic and/or intrinsic signaling that govern cellular turnover and organ size during development [59-61]. Lastly, clonal analysis and temporal recording applied to the *Apc*^*Min/+*^ mouse model functionally validated the possibility of polyancestral tumor initiation, to the extent that barcoded mutations can be traced back to multiple normal epithelial cell ancestors. The integrative analysis of the HTAN colorectal precancer atlas and mouse barcoding data allowed us to delineate factors that affect the earliest stages of tumor development, including clonal composition [107, 115, 116] and molecular signatures influencing the clonal fitness landscape [94, 105, 117]. Overall, our data suggests a continuum of selective pressures during tumorigenesis that modulates transition from a polyancestral composition in the early precancer stage to a monoancestral composition in the advanced cancer stage [20, 85, 94, 118]. In future, we may leverage evolutionary processes of human colorectal polyps to chart the multi-step progression of precancer to cancer that may illuminate strategies for early intervention.

Our multi-purpose platform is scalable and broadly applicable to a variety of studies beyond developmental recording. For example, in a companion paper (Islam et al.), we showed that NSC-seq enables single-cell CRISPR screens (Perturb-seq) using conventional vector libraries. This can be multiplexed with barcoding for simultaneous in vivo genetic perturbation and lineage tracking at single-cell resolution [119]. In addition, using biological signal-responsive Cas9 promoters [32], NSC-seq can be used for multifactorial recording of cellular and molecular activities [5, 7], as well as single-cell mapping of the neuronal connectome in the brain [120]. Overall, we envision that NSC-seq platform will expand the application of CRISPR technologies, and together, they will form powerful tools for the scientific community.

## Acknowledgement

This publication is part of the HTAN (Human Tumor Atlas Network) consortium paper package. The authors wish to thank the study participants and funding support by the HTAN grant U2CCA233291 (to R.J.C., K.S.L., and M.J.S.), TBEL U54CA274367 (to R.J.C., K.S.L., and M.J.S.), R35CA197570 and P50CA236733 (to R.J.C.), R01DK103831 (to K.S.L.), K07CA122451 (to M.J.S.). We thank members of the Lau and Coffey laboratories (especially Andrea Rolong and Matthew E. Bechard) for various assistances including animal housing, tissue imaging, and single-cell data collection. Cores used by this study included Survey and Biospecimen Shared Resource, TPSR (U24DK059637), VANTAGE, and REDCap (UL1TR000445). 1cellbio and RAN biotechnologies helped to synthesis the custom hydrogel beads. We also thank Alyssa Hasty (VU) and Angela Jones (VANTAGE) for their assistance. Vanderbilt University submitted a U.S. patent application for NSC-seq and M.I., R.J.C. and K.S.L. are listed as inventors. We use BioRender for drawing many schematics in this study. We apologize in advance to those we have failed to acknowledge due to space constraints.

## Authors contribution

Conceptualization, M.I. and K.S.L.; data curation, M.I., Y.Y., A.J.S., Y.X., P.M., M.A.R.-S., M.J.S., and K.S.L.; formal analysis, M.I., Y.Y., V.M.S., K.P.M., N.T., Z.C., M.A.R.-S., J.D., Q.L., and K.S.L.; investigation, M.I., D.W., I.S., I.J.M.,L.T.T, G.M.C., M.A.G., J.C.R., H.Z., K.C.; R.J.C., and K.S.L.; methodology, M.I., and K.S.L.; project administration, M.I.., A.J.S., M.J.S., R.J.C., and K.S.L.; resources, M.I., Q.L., M.J.S., R.J.C., and K.S.L.; software, M.I., Y.Y., K.P.M., and K.S.L.; supervision, M.I., G.M.C, M.A.G., I.J.M., K.C., H.Z., J.C.R., R.J.C., M.J.S., and K.S.L.; validation, M.I. and Y.Y., and K.S.L.; visualization, M.I. and Y.Y.; and K.S.L.; writing – original draft, M.I., R.J.C., and K.S.L.; writing – reviewing and editing, M.I., Y.Y., A.J.S., V.M.S., K.P.M., Y.X., N.T., Z.C., P.M., M.A.R.-S., I.S., J.D., K.C., M.A.M., J.C.R., I.G.M., D.W., Q.L., H.Z.,R.K., G.M.C., M.J.S., R.J.C., and K.S.L.

## Data and code accessibility

Human data have been deposited to the HTAN Data Coordinating Center Data Portal at the National Cancer Institute: https://data.humantumoratlas.org/ (under the HTAN Vanderbilt Atlas). Mouse data deposited in GEO: ^********^. Reviewer token:^*********^. NSC-seq data analysis pipeline reported in GitHub: https://github.com/Ken-Lau-Lab/NSC-seq.

## Supplemental Figures

Extended Data Fig. 1: Design and validation of NSC-seq platform.

Extended Data Fig. 2: Overview of temporal recording.

Extended Data Fig. 3: Mitochondrial variants detection and validation for lineage analysis. Extended Data Fig. 4: Cell-type annotation and data quality control metrics for mouse embryos.

Extended Data Fig. 5: Temporal recording reveals asymmetric contribution of early embryonic clones to germ layers and tissue types.

Extended Data Fig. 6: Catalog of cellular turnover across embryonic timepoints.

Extended Data Fig. 7: Lineage reconstruction of mouse embryogenesis.

Extended Data Fig. 8: Somite-derived hematopoiesis.

Extended Data Fig. 9: Gut endoderm development and progenitor specification.

Extended Data Fig. 10: Lineage convergence during gut endoderm development.

Extended Data Fig. 11: Clonal dynamics of adult intestinal epithelium.

Extended Data Fig. 12: Multi-omic analysis of human colorectal polyps.

Extended Data Fig. 13: Tracking clonal composition of murine intestinal adenomas.

